# Unbuttoning the impact of N501Y mutant RBD on viral entry mechanism: A computational insight

**DOI:** 10.1101/2020.12.30.424906

**Authors:** Tanuj Sharma, Mohammad Hassan Baig, Moniba Rahim, Jae June Dong, Jae-Yong Cho

## Abstract

The ongoing coronavirus disease 2019 (COVID-19) pandemic has become a serious global threat. Severe acute respiratory syndrome coronavirus-2 (SARS-CoV-2), the virus responsible for this pandemic has imposed a severe burden on the medical settings. The spike (S) protein of SARS-CoV-2 is an important structural protein playing a key role in the viral entry. This protein is responsible for the receptor recognition and cell membrane fusion process. The recent reports of the appearance and spread of new SARS-CoV-2 strain has raised alarms. It was reported that this new variant containing the prominent active site mutation in the RBD (N501Y) was rapidly spreading within the population. The reported N501Y mutation within the spike’s essential part, known as the ‘receptor-binding domain’ has raised several questions. Here in this study we have tried to explore the effect of N501Y mutation within the spike protein using several in silico approaches

## Background

Till now, three different human coronaviruses (CoVs) have been identified [1–3]. Severe acute respiratory syndrome (SARS-CoV), Middle East respiratory syndrome coronavirus (MERS-CoV), first reported in 2002 and 2012, were the first members of this pathogenic coronavirus family [1, 4, 5]. These pathogenic coronavirus family members were responsible for the outbreak in 2003 and 2012, respectively [1, 5, 6]. In December 2019, a new member of this family was identified, later named SARS-CoV-2, and soon became a ghost virus associated with the global pandemic [7]. Since the emergence of SARS-CoV-2, the most highly pathogenic coronavirus reported so far, its rapid spread all over the globe has posed a severe threat to public health. The pandemic caused by this new coronavirus has emerged as a significant threat and has severely affected almost a considerable world population as a whole. First time reported from the Wuhan city, China, in December 2019, SARS-CoV-2 has quickly spread worldwide. Among all outbreaks of the 21st century, the SARS-CoV-2 causing COVID-19 pandemic has the highest infection and death rate, 61 million infections, and over 1.4 million deaths on December 1, 2020.. The studies have found that the SARS-CoV-2 mediates its viral entry by binding to the human ACE2 receptor via its spike protein. The RBD region of spike protein is well reported to be involved in binding with the host ACE2. The overall binding mode of ACE2-SARS-CoV-2 RBD is almost similar to other counterpart, SARS-CoV, which also uses the similar viral entry mechanism [8–11]. All the RNA coronaviruses family members, SARS-CoV-2, CoV, and MERS, present an extensive rate of mutations [11–13]. These high mutation rates may be a factor accountable for these coronaviruses’ zoonotic nature and maybe a critical point leading the future risk of other members of this viral family to switch to humans from their traditional hosts. These mutations may develop resistance towards the antiviral drugs as well. The structural investigations identified that most of the SARS-CoV-2 RBD residues essential for its binding to ACE2 are highly conserved with those in the SARS-CoV RBD [14, 15]. A novel SARS-CoV-2 virus variant, referred to as SARS-CoV-2 VUI 202012/01 (Variant Under Investigation, year 2020, month 12, variant 01), has been recognized via viral genomic sequencing in the United Kingdom (UK). It is classified by means of numerous spike protein (protein responsible for the anchoring of the virus to the host cell) mutations (deletion 69-70, deletion 144, N501Y, A570D, D614G P681H, T716I, S982A, D1118H). The variant belongs to Nextstrain clade 20B [16], GISAID clade GR [3, 4], lineage B.1.1.7. Out of these mutations, the N501Y mutation modifies the spike’s essential part, known as the ‘receptor-binding domain’. This is where the spike makes initial contact with the body’s cell surface. Least changes in the structure make it uncomplicated for the virus to get within. N501Y was first sequenced in April 2020 in a virus in Brazil and is presently linked with a SARS-CoV-2 variant - an independent lineage from B.1.1.7, with an intensifying recurrence rate in South Africa [17]. Mutation N501Y is one of the six major contact residues present inside the receptor-binding domain (RBD) and has been acknowledged to increase the binding affinity to human cell-surface protein angiotensin converting enzyme 2 (ACE2) [18–20]. Changes in the receptor-binding region of the spike protein may ultimately lead to the change in the virus’s ACE2 binding specificity and alter antigenicity, i.e., recognition by immune antibodies the SARS-COV-2 virus becoming more contagious and spreading more effortlessly amid people. Here in this study, we have used Molecular dynamics and other computational approaches to explore the impact of this important point mutation on SARS-COV-2 RBD towards its binding against the receptor.

## Material and Methods

The structure of SARS-CoV-2 spike receptor-binding domain bound with ACE2 was retrieved from the protein data bank (6M0J) [21]. The Chain A of the complex was the Angiotensin-converting enzyme 2 (ACE2) while the chain E was the receptor-binding domain. Here we have mutated the Asn at 501 position of receptor-binding domain to Tyr. The mutant structure was energy minimized using both steepest descent minimization followed by the conjugate gradient minimization.

### Molecular Dynamics simulation

The wild type as well as the N501Y mutant of the complex of SARS-CoV-2 spike receptor-binding domain bound with ACE2 were subjected for MD simulation. The MD calculations was performed using CHARMM27 force field with GROMACS 20.2 package [22]. The system was initially solvated within the cubic solvation box with TIP3P water model and with 10 Å periodic boundary conditions. Further Na and Cl ions were added to satisfy the electro-neutrality condition. The system was subjected to energy minimization using the steepest descent integrator. The energy minimized model was subjected to NVT as well as NPT ensemble for 200 ps so as to stabilize the temperature and pressure respectively. The equilibrated system was further subjected to extended molecular dynamics simulation for 100 ns. Post molecular dynamics, analysis was done using chimera and grace.

## Results and Discussion

Declared as global pandemic by WHO in 11 March 2020, COVID-19 has become a serious threat to the worldwide population. While a large number of researches are being conducted, with several promising results providing a hope toward the treatment of COVID-19, still the threat is not seeming to be ending very soon. Today a large proportion of research investment has been made for the development of effective therapeutic candidate against SARS-CoV-2. After the announcement of several vaccines by different pharmaceutical groups, an optimistic speculation was made, but with the reports of new variants, this hopefulness provides an uncertainty and hints towards a long-term fight against the virus. The COG-UK consortium (https://www.cogconsortium.uk/about/) recently reported the appearance and spread of new SARS-CoV-2 strain. It was reported that this new variant containing the prominent active site mutation in the RBD (N501Y) was rapidly spreading within the population. Previous studies have reported the N501T mutation in the RBD has no prominent impact on its affinity toward ACE2 (when compare SARS-CoV-2 to SARS-CoV). But the impact of this newly reported mutant has not been investigated. Here, were performed the computational study to investigate the effect of this mutation on the binding affinity to the ACE2 and its impact on the transmission. The RBD domain of wild type strain of Covid19 has been explored and the structure of this SARS-CoV-2 spike receptor-binding domain bound with ACE2 protein has been reported[21]. The mutant N501Y was generated by built in rotamer generation Dunbrack 2010 library [23] in UCSF Chimera software. The complex generated by single mutation, was further subjected to 50000 steps of conjugate gradient minimization in presence of water so as to stabilize the complex to a more favorable conformation. In order to understand the molecular level interactions of the residue Asn501 with the ACE-2, we identified the residues occurring in the 5 Å radius. We observed that 4 residues of ACE-2, namely Gly352, Lys353, Asn355 and most importantly Tyr41 occurs in close proximity of 5 Å (figure-1B-D). Previous studies have well established the existence of cation-π interactions between cationic sidechain (Lys or Arg) and aromatic sidechain (Phe, Tyr, or Trp), when in close proximity [24–28]. The role of anion-π interaction (between an electron-deficient aromatic moiety and an anion conveniently located above the ring plane) has also been reported in many computational as well as experimental studies [29–32]. The geometry of the interaction indicates the involvement of a favorable cation-π or anion-π interaction. Careful investigation of the molecular orientation of these residues (around 501) indicated that the carboxamide group of Asn501 residue is perfectly oriented towards the 4-hydroxyphenyl group of the Tyr41. The orientation of the oxygen is particularly favoring the anion-π interaction with the π electron cloud of the phenyl ring. The distance observed between the oxygen atom and base of phenyl group was 2.924 Å, which is well within the required constraints of the distance between two atoms for the anion-π interactions (figure 1E). The distance observed between the Nitrogen atom and nearest atom of phenyl group was observed to be 5.295 Å, which indicates that formation of anion-π interaction might be more favored compared to cation-π interaction. Although, the formation of cation-π interaction cannot be ruled out. After the 100 ns MD run of both the wild type and mutated complex, we observed that the residue 501 plays a crucial role in interaction with the A chain of ACE-2 protein. Once the residue 501 is mutated to Tyr, we observed that there is a strong possibility of formation of double-sided anion-π interaction between the oxygen atom of each tyrosine with the π electron cloud of the phenyl ring of opposite Tyr residue. Also, there is possibility of π-π stacking interactions between the two aromatic phenyl rings of Tyr residue. In both of the above cases, the mutant strain would be observed to be more stably interacting with the ACE-2 protein than the wild type. During the course of the simulation, we observed that the π electron cloud of the phenyl ring of the Tyr501 acquired a perpendicular orientation which might result in the formation of T-shaped π-π stacking interaction between the Tyr501 of RBD domain and Tyr41 of ACE-2. Overall, the structure was observed to be highly stable and indicated a very low RMSD of less than 0.7 Å. Compared to mutant, wild type residue Asn501, indicated slightly higher RMSD of more than 1 Å, throughout the 100 ns MD run. Hydrogen bond analysis of the residue of the interest in the MD studies indicated that the mutant residue (Tyr501) was observed to form hydrogen bond interaction for a longer time than the Asn501 (figure-3). Similarly, RMSD plot indicated that the Tyr501 was observed to be more stable compared to the Asn501 (figure-4). Considering the poses observed in the mutant residue (Tyr501) and wild type (Asn501), it is evident that the mutant strain (N501Y) would interact with the ACE-2 protein with more stability and therefore might be more virulent that the wild type strain.

**Figure-1(A-E):**
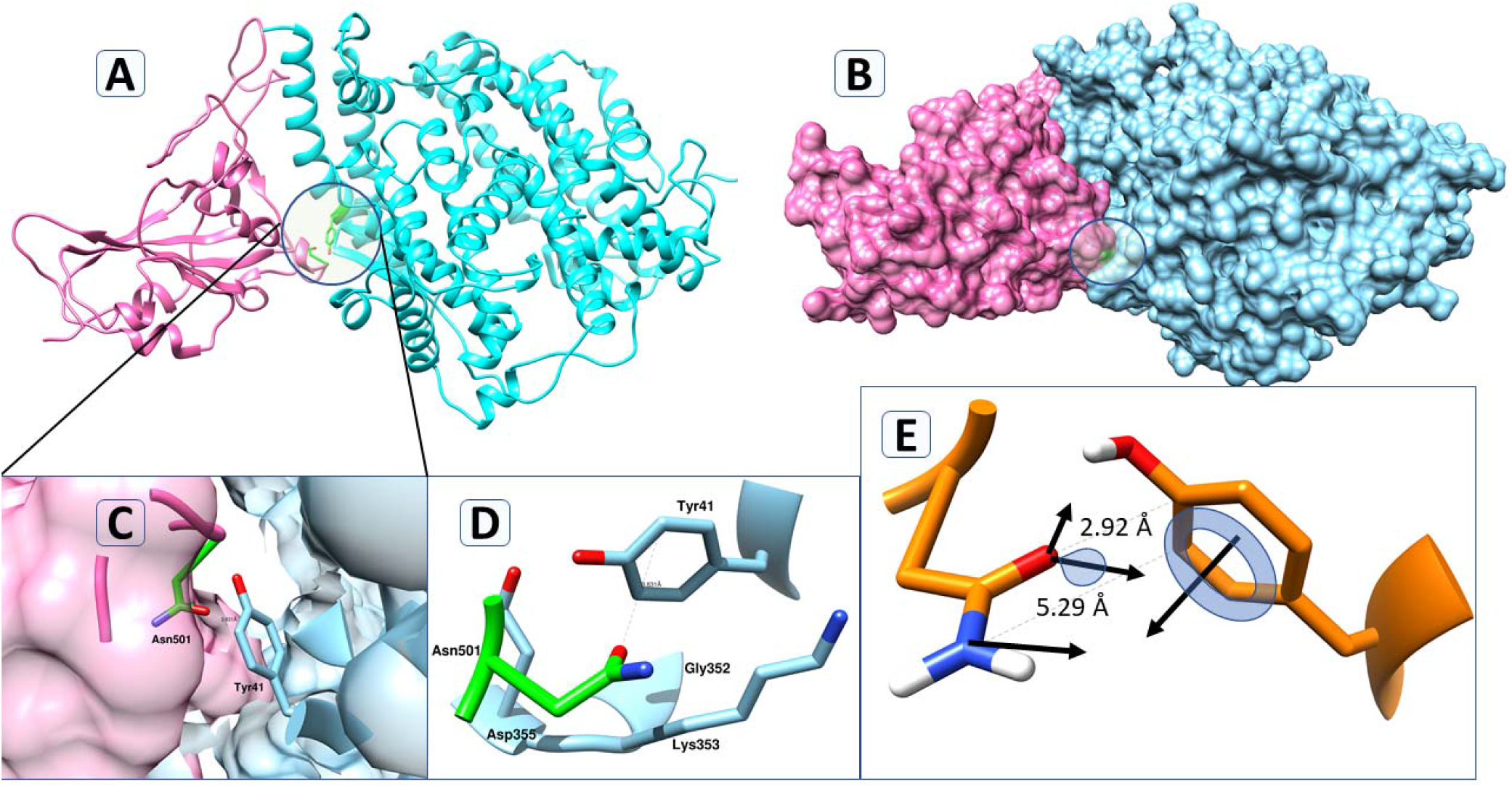
(A) The image indicates ribbon view model of the reported crystal structure of the wild type RBD domain with ACE-2 protein. (B) The image indicates the bound surface view model of the RBD domain (pink) of spike protein with the ACE-2 protein (blue). The slight green location within circle indicates the location of Asn501. (C) The image indicates the atom view model of the residue Asn501 with respect to Tyr41 of chain A of ACE-2. (D) The image indicates the residues observed with the 5 Å radius of the Asn501. (E) The image indicates the orientation of the Asn501 of RBD domain with respect to Tyr41 of ACE-2 protein. The lines indicate the vectors of electron clouds.

**Figure-2(A-G):**
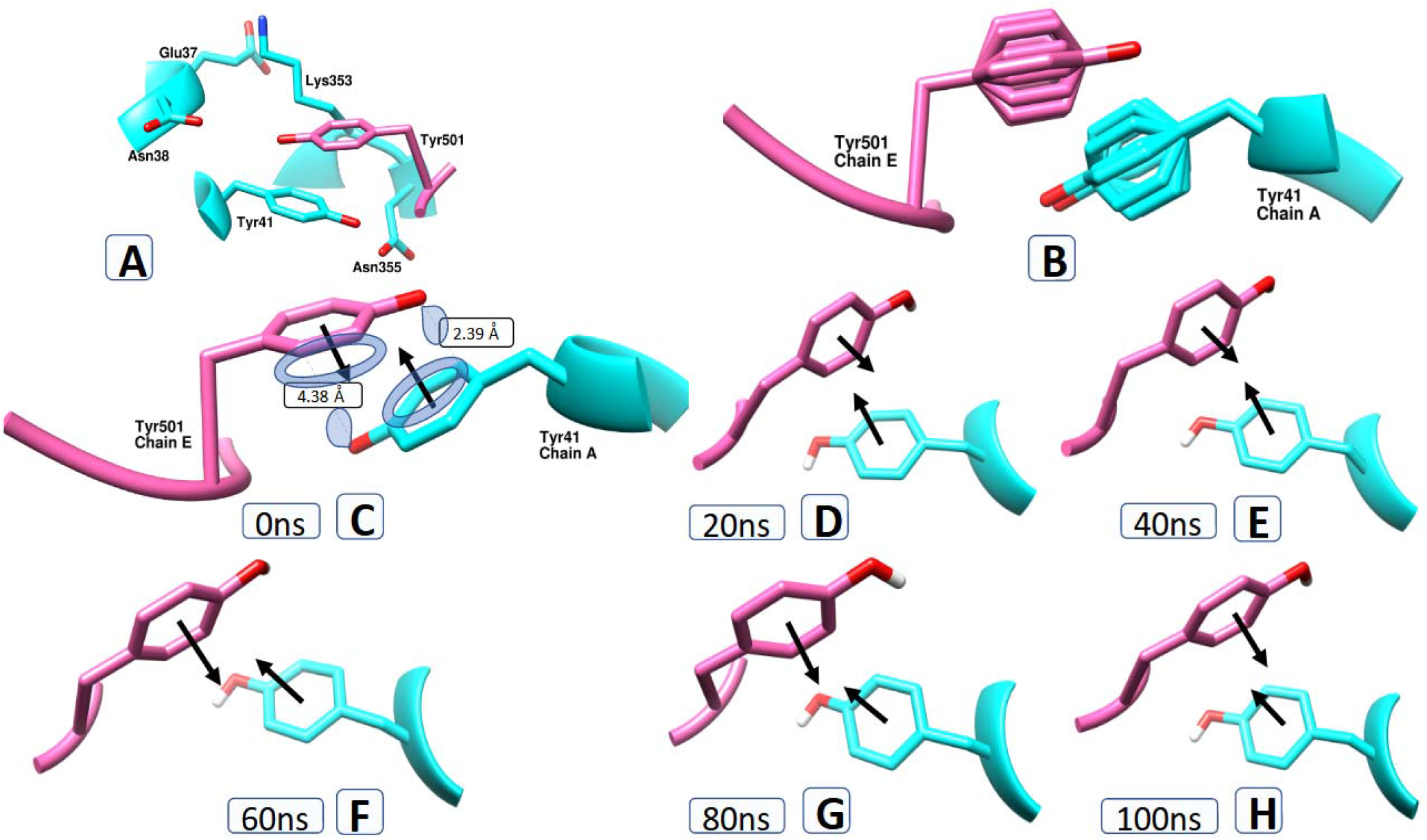
(A) The image indicates the residues occurring in 5 Å radius of Tyr501 residue. (B) The image indicates the different Tyr rotamers observed in both RBD domain of wild type as well as opposite Tyr41 position of ACE-2 protein. (C) The image indicates the orientation of Tyr501 of RBD and Tyr41 of ACE-2 rotamer having highest probability. The lines indicate the vectors of electron clouds of phenyl ring. (D-H) The image indicates the atom view model of the residue Tyr501 with respect to Tyr41 of chain A of ACE-2 at different time frames namely 20, 40, 60, 80 and 100 ns during MD.

**Figure-3:**
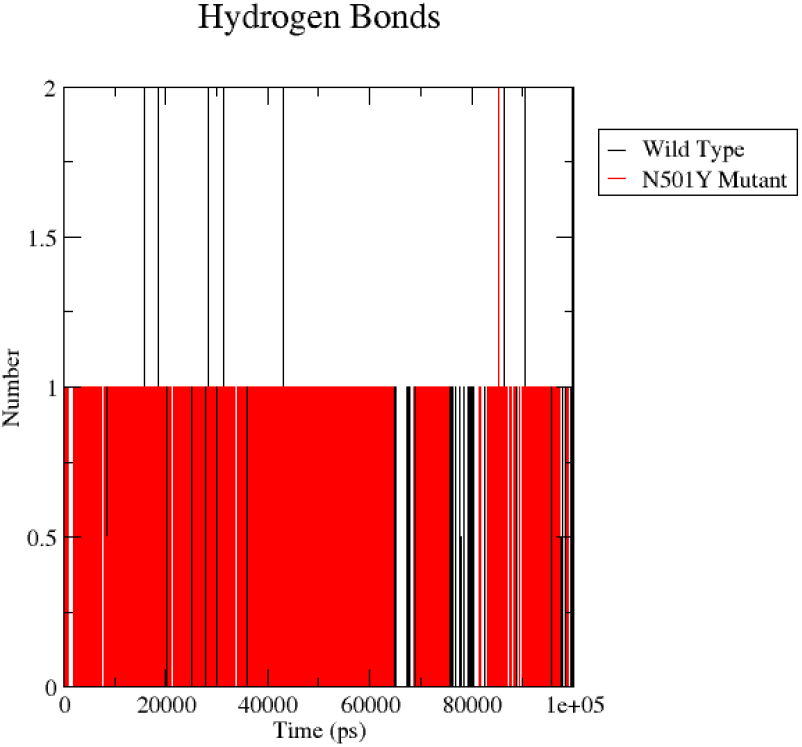
The image indicates the formation of hydrogen bonds between the residue at position 501 with the ACE-2 protein. The mutant type (Tyr501) residue was observed to be forming stable hydrogen bond with the A chain of ACE-2 protein than the wild type (Asn501).

**Figure-4:**
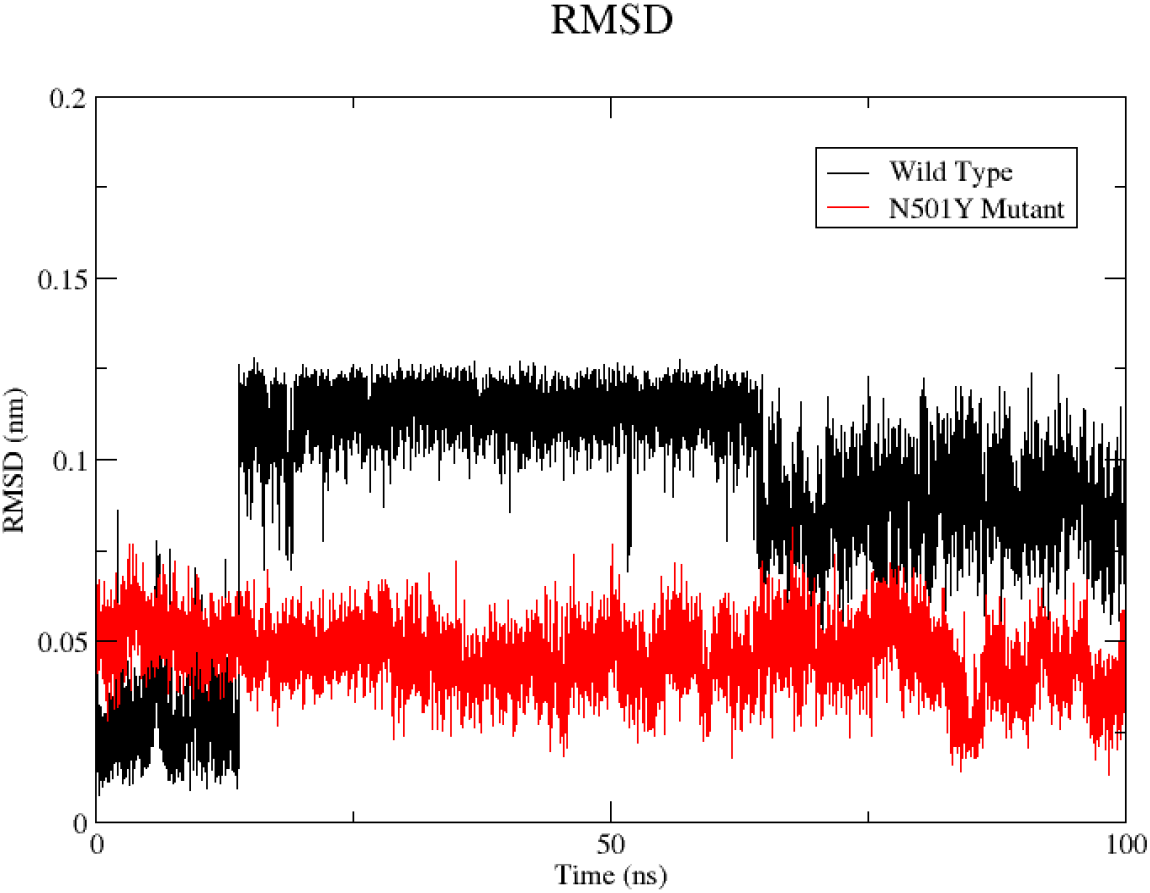
The image indicates the RMSD plot of the residue at position 501 in the RBD domain of the spike protein of the Covid19 protein. The mutant type (Tyr501) residue was observed to be more stable than wild type residue (Asn501) throughout the simulation indicating high stability of the mutant residue, in association with the ACE-2 protein.

## Conclusion

Coronavirus is a large family of viruses which have been around for a long duration. The human affecting SARS-CoV-2 which has caused havoc throughout the world probably has been infecting animals for quite some time. It’s new to the human, although not so considering the animal kingdom and is constantly evolving through undergoing mutations in its RNA. It is highly important to understand the various affects of the mutations on the protein expression, especially those proteins, which are involved in interactions with the human protein. The mutation N501Y in the RBD domain of the spike protein, which involves the residue, present on the interface of the spike protein and it is highly important to understand the behavior of this mutation on the interaction with the human ACE-2 protein. Through current computational studies, it is evident that the residue Asn501 as well as its mutant residue Tyr501 both plays crucial role in interaction with the ACE-2 protein. Most importantly the interactions with the Tyr41 of the human protein plays a crucial role for the stability of both wild type as well as mutant strain. Further, considering the molecular interactions observed during the MD studies, there is a strong possibility that the mutant N501Y might increase the stability of the RBD domain with the ACE-2 protein. This increase in stability of the RBD domain, may result in higher virulence of the mutant strain than its wild type strain. Further mutational studies on the interacting residues namely Tyr41, might give more clear understanding on the interaction. These interactions could also be used for the design and identification of vaccines and novel inhibitors against the newly emerging coronavirus strains.

## Notes

### Competing Interest Statement

The authors have declared no competing interest.

